# Coral *Symbiodinium* community composition across the Belize Mesoamerican Barrier Reef System is influenced by host species and thermal variability

**DOI:** 10.1101/154179

**Authors:** JH Baumann, SW Davies, HE Aichelman, KD Castillo

## Abstract

Reef-building corals maintain a symbiotic relationship with dinoflagellate algae of the genus *Symbiodinium* and this symbiosis is vital for the survival of the coral holobiont. *Symbiodinium* community composition within the coral host has been shown to influence a coral’s ability to resist and recover from stress. A multitude of stressors including ocean warming, ocean acidification, and eutrophication have been linked to global scale decline in coral health and cover in recent decades. Three distinct thermal regimes (high_TP_, mod_TP_, and low_TP_) following an inshore-offshore gradient of declining average temperatures and thermal variation were identified on the Belize Mesoamerican Barrier Reef System (MBRS). Quantitative metabarcoding of the ITS-2 locus was employed to investigate differences and similarities in *Symbiodinium* genetic diversity of the Caribbean corals *Siderastrea siderea*, *S. radians*, and *Pseudodiploria strigosa* between the three thermal regimes. A total of ten *Symbiodinium lineages* were identified across the three coral host species. *Siderastrea siderea* associated with distinct *Symbiodinium* communities, however *Symbiodinium* communities of its congener, *S. radians*, and *P. strigosa*, were more similar to one another. Thermal regime played a role in defining *Symbiodinium* communities in S. siderea but not *S. radians* or *P. strigosa*. Against expectations, *Symbiodinium trenchii*, a symbiont known to confer thermal tolerance, was dominant only in *S. siderea* at one sampled offshore site and was rare inshore, suggesting that coral thermal tolerance in more thermally variable inshore habitats is achieved through alternative mechanisms. Overall, thermal parameters alone were likely not the only primary drivers of *Symbiodinium* community composition, suggesting that environmental variables unrelated to temperature (i.e., light availability, or nutrients) may play key roles in structuring coral-algal communities in Belize and that the relative importance of these environmental variables may vary by coral host species.

## Introduction

Obligate symbioses, relationships in which two or more organisms depend on one another for nutrition and survival, occur globally. Such symbioses are ubiquitous in plants and algae, i.e., Mycorrhiza [1], lichens [2], or insects, i.e., ants and bacteria [3]. The effects of climate change are expected to disrupt proper functioning of many symbioses, including that of reef-building corals [4–6], who’s success depends on the symbiosis between the coral host and photosynthetic algae of the genus *Symbiodinium* [7–9]. Under stressful conditions this coral-*Symbiodinium* relationship breaks down, resulting in the loss of endosymbiont cells and/or photosynthetic pigments from the coral tissue in a process known as ‘coral bleaching’ [10]. Coral bleaching is most commonly associated with thermal stress [11–15] and is predicted to increase in frequency and severity as the world’s climate continues to change [5, 16–21]. Increased thermal stress resulting from climate change combined with other local stressors such as eutrophication, habitat destruction, and overfishing has created an uncertain future for coral reefs [6, 13, 22]. In the Caribbean Sea, warming rates are higher than in any other tropical basin [23, 24] and coral cover has declined by as much as 80% in recent decades [25]. It has been predicted that Caribbean coral reefs may suffer biannual bleaching events within the next 20-30 years [17] and annual bleaching by 2040 [26].

In the face of a changing climate and widespread reef declines, corals will need to rapidly increase their thermal tolerance in order to persist in their current form [18, 27]. Coral thermal tolerance has been shown to be influenced by a coral’s thermal history, which among other factors includes average environmental temperature and extent of thermal variability [28, 29]. On average, corals previously exposed to warmer temperatures show decreased mortality during bleaching events [30] and more stable growth patterns [31] compared with corals exposed to cooler temperatures, which exhibit greater mortality during heat stress and declining growth rates with increased temperatures [30, 31]. Exposure to greater daily thermal variation has also been shown to increase coral thermal tolerance [32] and has been associated with higher coral cover and slower mortality rates when compared to reefs exposed to less thermal variation [33]. Coral thermal tolerance is also heritable with larvae from parent colonies on lower-latitude (warmer) reefs showing a 10-fold increase in survival under heat stress when compared to larvae from cooler reefs locations [34]. A growing body of evidence suggests that the coral host plays a significant role in thermal tolerance [35–38], however, plasticity or specificity of coral-associated *Symbiodinium* and bacterial communities have also been shown to play a significant role in overall thermal tolerance [39–43].

The clades, lineages, or species of *Symbiodinium* hosted by a coral are critical to its survival and resilience to stress. The genus *Symbiodinium* is genetically diverse and comprises at least nine divergent clades [clades A-1; 44]. These clades can be further broken down into lineages, corresponding approximately to species level diversity [45], with some species conferring variable benefits [39, 44, 46]. In particular, some *Symbiodinium* are more thermally tolerant than others [9, 39, 47], specifically *Symbiodinium* clade D [48]. In contrast, clade C is more thermally sensitive [49–51], yet it includes *Symbiodinium thermophilum*, a thermally tolerant species within clade C endemic to the Red Sea [52]. This example illustrates that making clade level generalizations is problematic due to the physiological diversity within a single *Symbiodinium* clade [53]. Specific lineages within clades can also confer various advantages. For example, C1 enhances growth rate [54], *S. thermophilum* confers heat tolerance [52], and B2 confers cold tolerance [55]. Additionally, species D1a (*Symbiodinium trenchii)* has been shown to be both heat tolerant [56, 57], and cold tolerant [47]. However, the increased thermal tolerance of a coral which predominantly hosts clade D *Symbiodinium* appears to come at a cost of lower lipid stores, reproductive potential, growth, and carbon fixation rates compared with corals that host other clades [58–61]. Due to the high levels of variation in coral host-*Symbiodinium* interactions, it is essential to identify which lineages are present in order to help predict how a coral may respond to environmental stressors.

The majority of coral species host one dominant *Symbiodinium* lineage [44, 62, 63] along with several non-dominant lineages [64], each proliferating primarily by asexual cloning [53]. However, other corals can host multiple dominant lineages or clades [39, 53]. Recent advances in genetic techniques, especially next-generation sequencing (NGS), have allowed researchers to identify cryptic and low-abundance symbionts comprising 0.1% or more of the total *Symbiodinium* community within a host [37, 65]. It is important to understand these low-abundance *Symbiodinium*, as they have the potential to play important roles in coral-algal holobiont physiology under ambient and stressful conditions [66–68, but see also 69]. Identifying trends in *Symbiodinium* community variation (including cryptic or low abundance lineages) within and between species across a coral reef may allow for a better understanding of the role of *Symbiodinium* communities in modulating coral response to environmental variation.

*Symbiodinium* communities have been shown to vary regionally [between reef systems; 61, 70, 71], locally [within a reef system; 70], temporally [across time on the same reef; 72], and within a colony [71]. Studies of this variation have revealed geographically endemic lineages of *Symbiodinium* which may play a significant role in local and regional scale coral survival and stress tolerance [39, 71, 73]. While temperature stress may play a role in structuring *Symbiodinium* communities [74], variations in other environmental factors have also been shown to drive *Symbiodinium* community composition. For example, physical processes and total suspended solids (a proxy for nutrients and flow) drive *Symbiodinium* associations within the *Orbicella annularis* species complex in Belize and Panama [70]; however, on a regional scale (e.g., the entire Caribbean Sea), *O. annularis Symbiodinium* communities differed based on patterns of chronic thermal stress [75]. Additionally, the presence of several subclades of *Symbiodinium* correlated with other environmental parameters, such as cooler summers, nutrient loading, and turbidity [75]. Taken together, these studies demonstrate that variation in *Symbiodinium* communities can be driven by a variety of environmental parameters and may be specific to each coral species in each specific environment.

The majority of Caribbean *Symbiodinium* biogeography studies have focused on the *Orbicella* species complex [70, 71, 75] as *Orbicella* spp. has experienced significant declines over the last two decades [76] and are now listed as ‘threatened’ under the Endangered Species Act. However, the variation in *Symbiodinium* communities of other more stress tolerant corals, such as *Sidereastrea siderea* and *S. radians* [77–82], remain relatively understudied. Here, we assess *Symbiodinium* community composition in three species of ubiquitous Caribbean corals (*Siderastrea siderea*, *S. radians*, and *Pseudodiploria strigosa*) across three distinct thermal regimes along the Belize Mesoamerican Barrier Reef System (MBRS) previously shown to influence coral community composition [83]. Coral-associated *Symbiodinium* communities were examined across an inshore-offshore thermal gradient and a latitudinal gradient to elucidate the role that coral species, local habitat, and thermal regime play in structuring *Symbiodinium* communities in the western Caribbean Sea.

## Methods

### Site selection and characteristics

Ten sites along the Belize MBRS were selected. These sites were previously characterized into three thermally distinct regimes (low_TP_, mod_TP_, high_TP_) and exhibited variations in coral species diversity and richness [83]. High_TP_ sites (inshore) were characterized by larger annual temperature variation, higher annual maximum temperatures, and are exposed to temperatures above the regional bleaching threshold of 29.7°C (Aronson et al., 2002) more often than mod_TP_ sites (mid-channel reefs) and low_TP_ sites (offshore) [83]. High_TP_ sites were dominated by stress tolerant and weedy coral species while corals representing all four coral life histories [stress tolerant, weedy, competitive, and generalist; 82] were present in low_TP_ and mod_TP_ sites [83].

### Sample Collection

In November 2014, five to ten (quantity depended on local availability) coral tissue microsamples (approx. 2 mm diameter) were collected at 3 to 5 m depth from three coral species (*Siderastrea siderea*, *S. radians*, and *Pseudodiploria strigosa*) at nine sites across four latitudes along the Belize MBRS (Fig 1; Table 1). Each latitudinal transect contained a low_TP_, mod_TP_, and high_TP_ site. The transects from north to south were: Belize City, Dangriga, Placencia, and Punta Gorda (Fig 1). All three sites within the Punta Gorda and Placencia transects were sampled, but only the low_TP_ and high_TP_ sites were sampled along the Belize City and Dangriga transects due to time constraints. Samples collected at the Belize City high_TP_ site were collected in October 2015, as no corals were located in the area in 2014, but patch reefs were located in 2015. Coral microsamples were collected at least 1m apart from one another to randomize micro-environmental and host genetic effects in order to attain more site-specific representative samples. Microsamples were collected from colony edges to avoid unnecessary damage to the larger colony and to limit effects of *Symbiodinium* zonation within an individual [71]. Tissue microsamples were placed on ice immediately following collection for transport to mainland Belize. Microsamples were then preserved in 96% ethanol and stored on ice at −20°C, and transported on ice to the coral ecophysiology lab at the University of North Carolina at Chapel Hill and stored at −20°C until DNA isolation.

**Fig 1:**
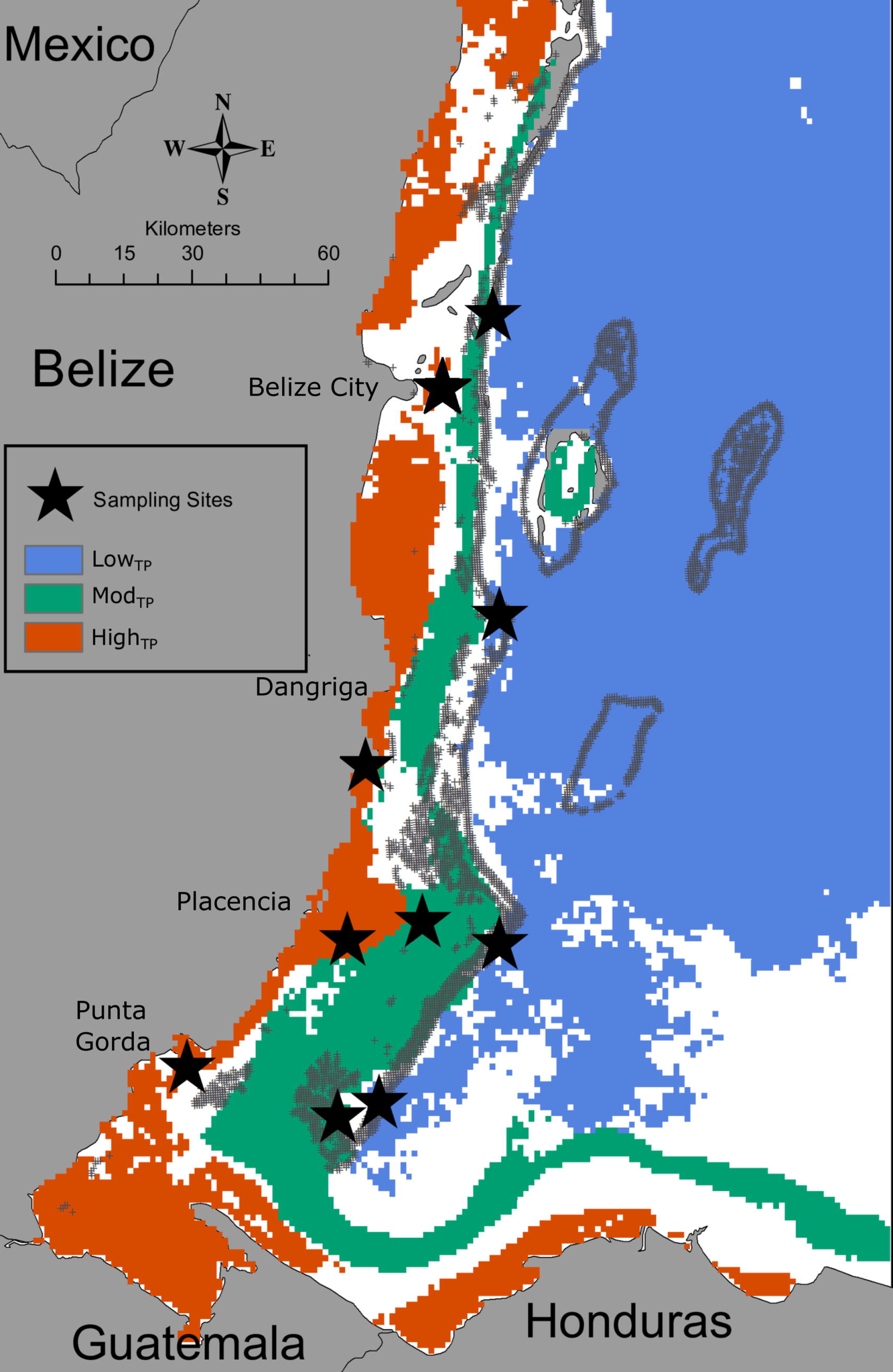
Thermal regime designations for sampling sites on the Belize MBRS [83]. Stars indicate sites where coral tissue samples were collected for *Symbiodinium* community analysis. Low_TP_, mod_TP_, and high_TP_ are defined based on combined averages of annual maximum temperature, annual temperaturerange, annual days above the bleaching threshold, and annual longest streak of consecutive days above the bleaching threshold. LOWTP sites exhibit the lowest values for all parameters measured and high-ip sites exhibit the highest. A more detailed description of classification of these thermal regimes can be found in Baumann et al. [83].

**Table 1:** Sampling locations and sample size for *S. siderea* (SSID), *S. radians* (SRAD), and *P. strigosa* (PSTR). Locations are listed in order of descending latitude (Northernmost to Southernmost). ‘-‘ represent an instance where sample size is equal to zero (n=0).

| Transect | Thermal<br>regime | Collection<br>Date | Illumina<br>Lane | Lat (°N) | Long<br>(°W) | SSID | SRAD | PSTR |
| --- | --- | --- | --- | --- | --- | --- | --- | --- |
| Belize City | Low | Nov 2014 | 2 | 17.64363 | 88.0264 | n=10 | --- | --- |
| Belize City | High | Oct 2015 | 2 | 17.48685 | 88.1207 | n=10 | --- | --- |
| Dangriga | Low | Nov 2014 | 2 | 17.078 | 88.01285 | n=9 | --- | --- |
| Dangriga | High | Nov 2014 | 2 | 16.79491 | 88.27699 | n=10 | --- | --- |
| Placencia | Low | Nov 2014 | 1 | 16.45816 | 88.01295 | n=7 | n=7 | n=5 |
| Placencia | Mod | Nov 2014 | 1 | 16.49995 | 88.16527 | n=6 | n=7 | n=6 |
| Placencia | High | Nov 2014 | 1 | 16.4654 | 88.31315 | n=9 | n=9 | n=5 |
| Sapodilla | Low | Nov 2014 | 1 | 16.15729 | 88.25073 | n=8 | --- | --- |
| Sapodilla | Mod | Nov 2014 | 1 | 16.13013 | 88.33234 | n=6 | --- | n=6 |
| Sapodilla | High | Nov 2014 | 1 | 16.2245 | 88.62943 | n=8 | n=6 | --- |

### Sea Surface Temperature

Daily 1-km horizontal resolution sea surface temperature (SST) estimates were acquired from the NASA Jet Propulsion Laboratory’s Multi-Scale High Resolution SST (JPL MUR SST) product via NOAA Environmental Research Division’s Data Access Program (ERDDAP-https://coastwatch.pfeg.noaa.gov/erddap/index.html) [84] and analyzed following Baumann et al [83]. Several additional temperature parameters were taken into account for this study, including: annual degree heating days (similar to degree heating weeks, as per Gleeson and Strong [85]), annual minimum temperature, annual average temperature, annual winter average temperature, and annual summer average temperature. Values for these parameters within the three thermal regimes are reported in Table S1.

### DNA Extraction

Coral holobiont (coral, algae, and microbiome) DNA was isolated from each sample following a modified phenol-chloroform [86–88] method described in detail by Davies et al (2013). Briefly, DNA was isolated by immersing the tissue in digest buffer (100 mM NaCL, 10mM Tris-Cl pH 8.0, 25 mM EDTA pH 9.0, 0.5% SDS, 0.1 mgml^-1^ Proteinase K, and 1 μgml^-1^ RNaseA) for 1 h at 42°C followed by a standard phenol-chloroform extraction. Extracted DNA was confirmed on an agarose gel and quantified using a Nanodrop 2000 Spectrophotometer (Thermo Scientific).

### PCR amplification and metabarcoding

The ITS-2 region (350 bp) was targeted and amplified in each sample using custom primers that incorporated *Symbiodinium* specific ITS-2-dino-forward and its2rev2-reverse regions [65, 73, 89]. Each primer was constructed with a universal linker, which allowed for the downstream incorporation of Illumina specific adapters and barcodes during the second PCR as well as four degenerative bases whose function was to increase the complexity of library composition. The forward primer was 5’-<u>GTCTCGTCGGCTCGG</u> + *AGATGTGTATAAGAGACAG* + NNNN + **CCTCCGCTTACTTATATGCTT**-3’ where the underlined bases are the 5’-universal linker, italicized bases indicate spacer sequences, N’s denote degenerative bases and the bold bases are the ITS-2-dino. The reverse primer was 5’-<u>TCGTCGGCAGCGTCA</u> + *AGATGTGTATAAGAGACAG* + NNNN + **GTGAATTGCAGAACTCGTG**-3’.

Each 20uL PCR reaction contained 5-100 ng DNA template, 12.4 μL MilliQ H_2_O, 0.2 μM dNTPs, 1μM forward and 1μM reverse primers, 1X *Extaq* buffer, and 0.5 U (units) *Extaq* polymerase (Takara Biotechnology). PCR cycles were run for all samples using the following PCR profile: 95°C for 5 min, 95°C for 40 s, 59°C for 2 min, 72°C for 1 min per cycle and a final elongation step of 72°C for 7 min. The optimal number of PCR cycles for each sample was determined from visualization of a faint band on a 2% agarose gel (usually between 22 and 28 cycles) as per Quigley et al. (2014). PCR products were cleaned using GeneJET PCR purification kits (Fermentas Life Sciences) and then a second PCR reaction was performed to incorporate custom barcode-primer sequences [65] modified for Illumina Miseq as in Klepac et al. [90]. Custom barcode primer sequences included 5’-*Illumina adaptor* + 6 bp **barcode sequence** + <u>one of two universal linkers</u>-3’ (e.g.: 5’- *CAAGCAGAAGACGGCATACGAGAT* + **GTATAG** + <u>GTCTCGTGGGCTCGG</u>-3’, or 5’-*AATGATACGGCGACCACCGAGATCTACAC* + **AGTCAA** + <u>TCGTCGGCAGCGTC-3’</u>). Following barcoding, PCR samples were visualized on a 2% agarose gel and pooled based on band intensity (to ensure equal contributions of each sample in the pool). The resulting pool was run on a 1% SYBR Green (Invitrogen) stained gel for 60 minutes at 90 volts and 120 mAmps. The target band was excised, soaked in 30 uL of milli-Q water overnight at 4°C, and the supernatant was submitted for sequencing to the University of North Carolina at Chapel Hill High Throughput Sequencing Facility across two lanes of Illumina MiSeq (one 2×250, one 2×300). The two lanes produced similar mapping efficiencies (73% and 73%, respectively; Table S3).

### Bioinformatic Pipeline

The bioinformatic pipeline used here builds upon previous work by Quigley et al. [65] and Green et al. [73]. Raw sequences were renamed to retain sample information and then all forward (R1) and reverse (R2) sequences were concatenated into two files, which were processed using CD-HIT-OTU[91]. CD-HIT-OTU clusters concatenated reads into identical groups at 100% similarity for identification of operational taxonomic units (OTUs). Each sample was then mapped back to the resulting reference OTUs and an abundance count for each sample across all OTUs was produced. A BLASTn search of each reference OTU was then run against the GenBank (NCBI) nucleotide reference collection using the representative sequence from each OTU to identify which *Symbiodinium* lineage was represented by each OTU (Table S2).

The phylogeny of representative sequences of each distinct *Symbiodinium* OTU was constructed using the PhyML tool [92, 93] within Geneious version 10.0.5 (http://geneious.com) [94]. PhyML was run using the GTR+I model (chosen based on delta AIC values produced from jModelTest [92, 95]) to determine the maximum likelihood tree. The TreeDyn tool in Phylogeny.fr was used to view the tree (Fig 2) [96–98]. The reference sequences included in the phylogeny were accessed from GenBank (Table S6).

**Fig 2:**
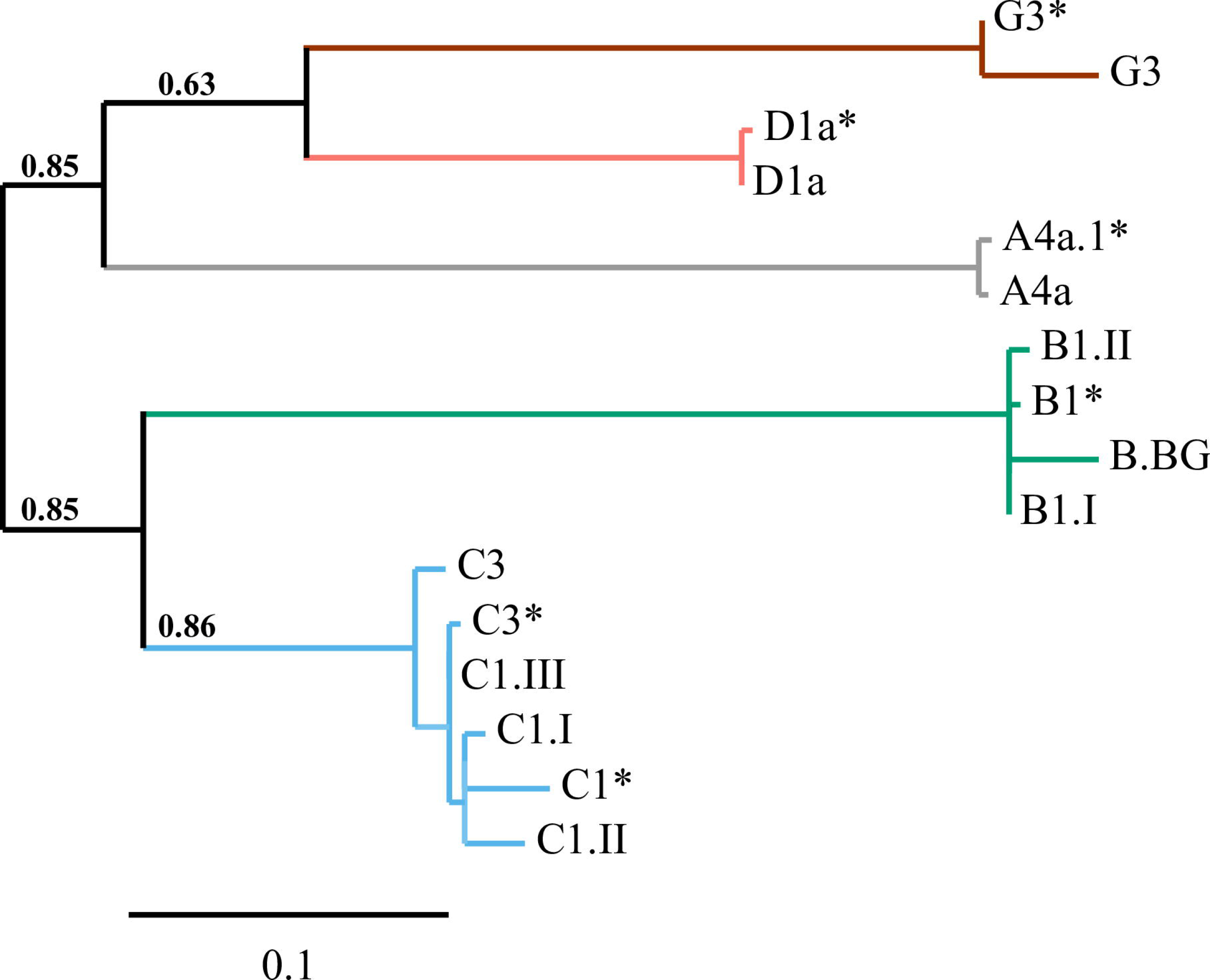
Phylogenetic analysis of ITS-2 sequences of representative OTUs from this study in addition to reference sequences for each clade (indicated by *). Branch support values are shown on the branches at divisions between distinct clades. The scale bar represents replacements per nucleotide site.

### Statistical Analysis

OTU abundance analysis used the R [99] package *MCMC*.*OTU* and followed methods described in Green et al. [73]. First, outlier samples with low sequence coverage (total log counts ≥2.5 standard deviations below the mean of all samples) were identified and removed, which removed 3 samples. Next, rare OTUs (<0.1% of the global sum of counts [as per 65]) were identified and discarded leaving 56 of the original 5,132 OTUs. Many remaining OTUs were identified as having the same *Symbiodinium* lineage (i.e., C1 or D1a) and these OTUs were regressed against one another. Positive correlations between OTUs within a lineage may indicate paralogous loci from the same genome [37, 73]. As a result, reads from OTUs within the same lineage that showed a positive R^2^ and significant *p*-value following linear regression were pooled in order to control for possible overestimation of biodiversity [100]. Pooling resulted in a final OTU table containing ten OTUs (Table S2). Raw reads, trimmed reads, mapped reads, and percentage of reads mapped per species were calculated and reported in Table 2. Final pooled OTUs were run through the MCMC.OTU package in R and fit to a model that included fixed effect for host species, collection site, and thermal regime (Table S4). Differences between fixed effects were calculated based on their sampled posterior distributions and statistical significance was calculated as per Matz et al. [101]. OTU count data were converted to relative abundances (%), which were used to generate Fig 3 (Table S5).

**Table 2:** Average number of raw reads, trimmed reads, and mapped reads including mapping efficiency (% of trimmed reads that mapped) for each species.

| Species | Raw reads | Trimmed reads | Mapped reads | Mapping<br>efficiency |
| --- | --- | --- | --- | --- |
| *S. siderea* | 46161 | 28453 | 22048 | 73% |
| *S. radians* | 51081 | 46812 | 35290 | 75% |
| *P. strigosa* | 88888 | 43928 | 31873 | 69% |
| **Total** | 186130 | 118834 | 89211 | 75% |

**Fig 3.**
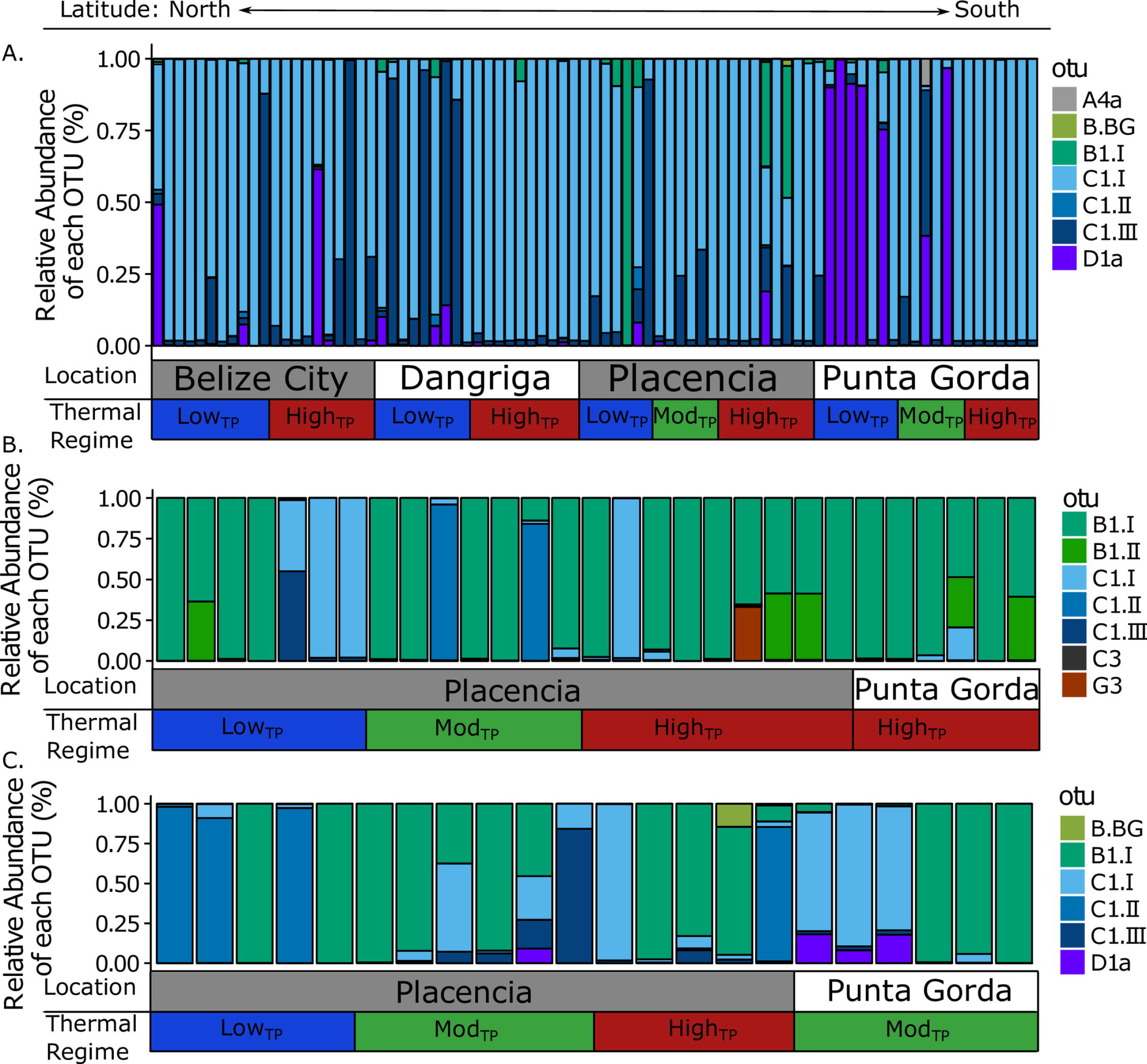
Relative abundance (%) of each OTU (lineage) in *S. siderea* (A), *S. radians* (B), and *P. strigosa* (C). Each column represents an individual sample. Columns are arranged by latitudinal transect (as indicated by site names in alternating gray and white boxes) and then by thermal regime (blue boxes indicate low_TP_ sites, green boxes indicate mod_TP_ sites, and red boxes indicates high_TP_ sites.

To visualize differences in symbiont communities between temperature regimes, latitude, and species, principal component analyses (PCA) were performed on all OTUs that passed filtering using the *vegan* package in R [102]. Count data were transformed using Bray-Curtis similarity and were used as input for PCA. PERMANOVA was carried out on each PCA using the *adonis* function of the *vegan* package in R [102].

## Results

### Symbiodinium diversity and abundance across the Belize MBRS

Our analysis produced 118,834 unique sequences of which 89,211 mapped to 10 OTUs (Table 1). The dominant OTU (hereafter referred to as lineage) in *S. siderea* was C1.I (74.39%), while B1.I dominated *S. radians* (70.31%) and *P. strigosa* (51.74%) samples (Table S5, Fig 3). Nine out of ten *Symbiodinium* lineages were present in *S. siderea* and *P. strigosa* while all ten were present in *S. radians* (Table S5). The four most abundant lineages in *S. siderea* were C1.I, C1.III, D1a, and B1.I (74.39%, 12.94%, 9.29%, and 2.94%, respectively; Table S5, Fig 3A) and date of collection did not impact the dominate *Symbiodinium* lineages (all samples collected in 2014 except for Belize City high_TP_ which were collected in 2015; Fig 3). *Symbiodinium* D1a (*S. trenchii*) was most abundant in *S. siderea* at low_TP_ sites, particularly the low_TP_ site along the most southern Punta Gorda transect (Table S5, Fig 3A) and lineage C1.III was more abundant in central and northern Belize (Belize City and Dangriga transects) compared to southern Belize (Figs 1, 3). Lineages C1.II, B1.II, G3, A4a, and B.BG were also present in *S. siderea* (Table S5, Fig 3A).

The four most abundant lineages in *S. radians* were B1.I, C1.I, B1.II, and C1.II (70.31%, 13.41%, 6.54%, and 2.19% respectively; Table S5, Fig 3B). B1.I was the dominant symbiont across all thermal regimes and all latitudes, but C1.I and C1.II were the most abundant *Symbiodinium* lineages in several samples from the central Placencia transect (Table S5, Fig 3B). Lineage C1.II was only present in proportions above 1% in 2 samples, both from the mod_TP_ site along the Placencia transect (Table S5, Fig 3B). D1a (*S. trenchii*) was only present in low abundance in *S. radians* (Table S5, Fig 3B). Lineages C1.III, D1a, G3, A4a, B.BG, and C3 were also present in *S. radians* (Table S5, Fig 3B).

The four most abundant lineages in *P. strigosa* were B1.I, C1.I, C1.II, and C1.III (51.74%, 21.87%, 16.92%, and 6.24%, respectively). C1.II was the most abundant lineage at the low_TP_ site in the Placencia transect, but B1.I was most abundant at all other sites (Table S5, Fig 3). C1.I was the second most abundant lineage in mod_TP_ and high_TP_ sites and C1.II was the second most abundant lineage in the low_TP_ site (Table S5, Fig 3C). D1a (*S. trenchii*) was only present in low abundance in *P. strigosa* (Table S5, Fig 3C). Lineages D1a, B1.II, G3, A4a, and B.BG were also present in *P. strigosa* (Table S5, Fig 3C).

### Host species specificity in <u>Symbiodinium</u> community composition

*Symbiodinium* communities differed significantly between *S. siderea* and the other two coral host species (Table S4, Fig 4A, *p*-value=0.001). This difference appears to be driven by higher relative abundances of C1.I and D1a (*S. trenchii*) in *S. siderea* compared to *P. strigosa* and *S. radians* (Fig 3A). Within *S. siderea*, *Symbiodinium* communities varied by thermal regime site, and latitude (Table S4, Fig 4B). *Symbiodinium* communities in *S. radians* and *P. strigosa* did not differ significantly by thermal regime, site, or latitude (Table S4).

**Fig 4.**
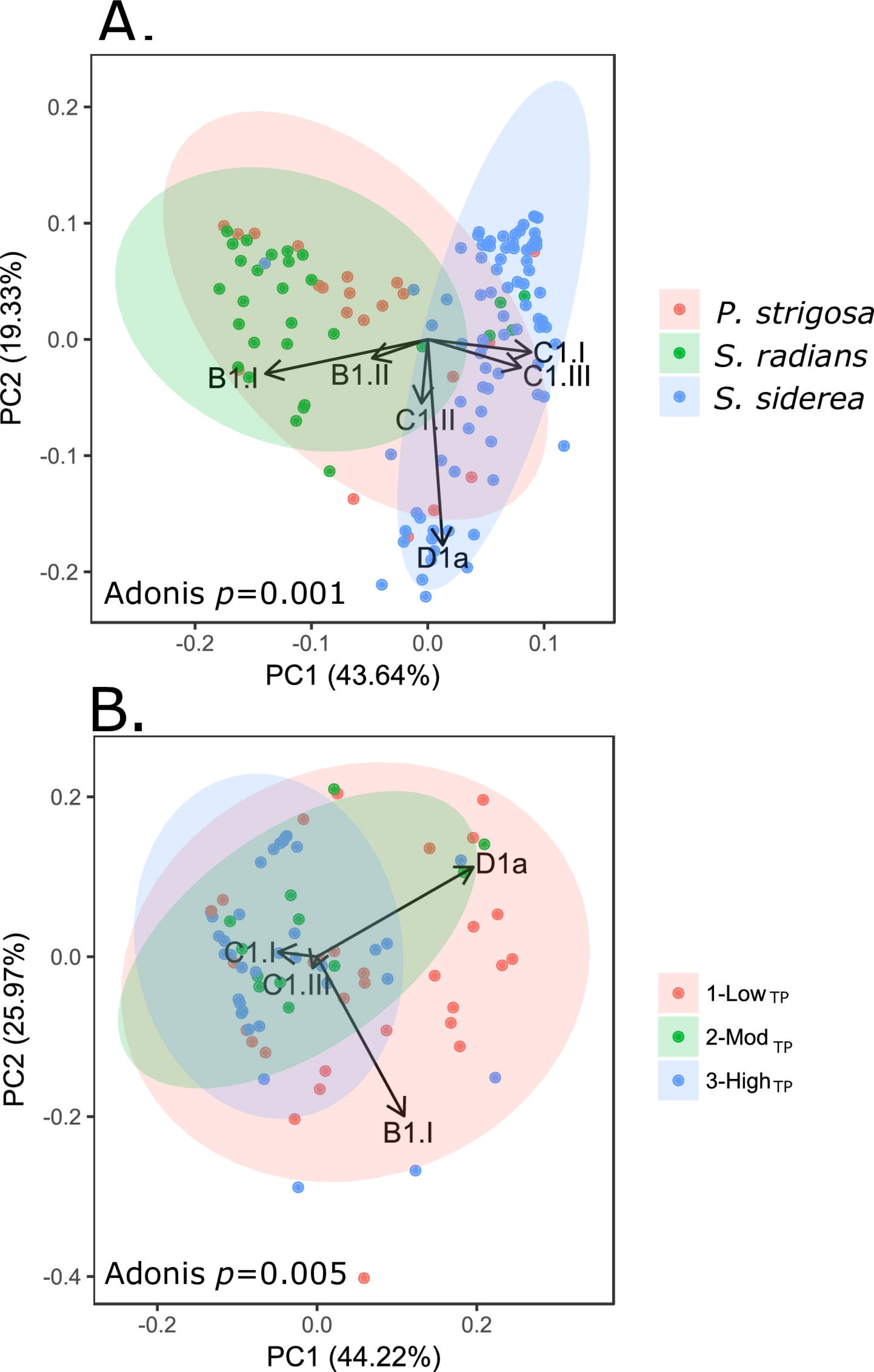
Principal component analysis (PCA) plots of *Symbiodinium* communities by species (A) and by thermal regime for *S. siderea* (B). Percentages on each axis indicate the amount of variation explained by each axis. Adonis *p*-*values* indicate significant results of PERMANOVA tests. See Table S4 for additional PERMANOVA results. Black arrows indicate loadings showing the magnitude and direction of the effect of each OTU on the total variance. Colored ellipses indicate 95% confidence intervals.

## Discussion

### Host-specificity drives <u>Symbiodinium</u> community composition

This study indicates that *Siderastrea siderea* hosts significantly different *Symbiodinium* communities from *S. radians* and *P. strigosa* on the Belize MBRS (Table S5, Fig 3), providing evidence to support previous findings of high rates of host-specific *Symbiodinium* associations within the Caribbean Sea where at least 62 genetically different *Symbiodinium* have been found and where >50% of *Symbiodinium* lineages have been found in only one coral genus [53, 103]. This trend contrasts that of the Indo-Pacific where *Symbiodinium* diversity is lower and a few host-generalist *Symbiodinium* associate with many corals [103]. The three coral species studied here were found to be associated with the two most abundant *Symbiodinium* clades in the Caribbean [104]: B1 in *S. radians* and *P. strigosa* colonies and C1 in *S. siderea* (Table S5, Fig 3). These associations are consistent with previous studies that identified the same dominant *Symbiodinium* in these species on the Belize MBRS [103]. However, our data contrast with findings of other studies on the same species elsewhere in the Caribbean which have identified other dominant *Symbiodinium* lineages in these host species [e.g., C3 and B5a in S. siderea and B5 and C46a in S. radians; 103, 105]. This supports previous evidence for regional endemism within the Caribbean Sea [103, 106]. *Symbiodinium* clade G, a lineage found in Octocorals [107], Foraminifera [108], and Pacific Porites spp. [109], was also observed to be a minor player in the symbiont communities of *S. radians* and *P. strigosa* (Table S5, Fig 3). This results indicates that this clade is present in the Caribbean Sea, however because this clade is not traditionally associated with Scleractinian corals, we cannot be confident that its presence is as a symbiont, a contaminant from the local environment, or that it was ingested as food. Differences in *Symbiodinium* communities between coral host species appear to be driven by the relative abundance of B1 and C1 as well as the presence or absence of D1a (Fig 4A). Presence of multiple lineages of C1 and B1 in this study (Table S2, Table S5) support previous evidence of phylogenetic partitioning, or highly specific lineages, in clades B and C [71, 103, 110, 111]. Interestingly, *Symbiodinium* communities were more similar between *S. radians* and *P. strigosa* than between *S. radians* and *S. siderea*, indicating that members of the same coral genus do not necessarily share a common dominant *Symbiodinium* partner, a phenomenon previously observed in *Siderastrea spp*. and *Orbicella spp*. across the Caribbean Sea [103]. Finney et al [103] show that *S. radians* and *S. siderea* exhibit different dominant *Symbiodinium* in both Belize (B5 vs. C1) and Barbados (B5 vs. C3). A similar trend is seen in *O. faveolata* and *O. annularis* (B17 vs. D1a in Belize and C7 vs. B1 in Barbados) [103]. These results suggest that *Symbiodinium* communities may not be influenced by coral host genus. Previously, it has been shown that symbiont acquisition strategy does not play a large role in determining *Symbiodinium* communities, however geographic distance and differences in environmental variables between habitats have been proposed as possible drivers of symbiont community composition [53, 103]. Coral life history strategy [82] or energetic demands may also play a role. Future research is needed to better understand this process. Differences in *Symbiodinium* communities between *S. siderea* and *S.* radians/ P. *strigosa* is suggestive that corals species are differentially affected by the environmental gradients sampled here.

### Thermal regime affects Symbiodinium community composition in S. siderea, but has no effect on other species

*Symbiodinium* communities varied significantly across thermal regimes in *S. siderea* (Table S4, Fig 4B), supporting previous evidence that habitat type [112] and temperature [113] are correlated with differences in *Symbiodinium* associations. *Symbiodinium* communities did not differ significantly across thermal regimes in *S. radians* or *P. strigosa*, possibly due to low sample size at each sampling site for these two coral species (Table 1; Fig 3). *Symbiodinium* communities did not differ between thermal regimes in *S. radians* or *P. strigosa* (Table S4), In this study, only temperature parameters were quantified, yet it is likely that they did not account for all of the variance in *Symbiodinium* communities for any coral host species investigated as other local impacts, such as nutrients, light availability, and/or sedimentation may play a role [48, 114–118].

### Role of local impacts on Symbiodinium communities

It has previously been shown that prevalence of specific *Symbiodinium* types within a coral host species can differ based on local scale environmental parameters such as nutrient loading and turbidity [75]. While these variables were not quantified in this study, chlorophyll-a ( *chl-a*), a proxy for nutrient input, has previously been shown to be positively correlated with thermal regime in Belize. Specifically, high_Tp_ sites had higher *chl-a* than low_Tp_ sites across the Belize MBRS [83]. Therefore, a PERMANOVA that shows significant differences in *Symbiodinium* communities between thermal regimes includes a confounding effect of nutrient input (Table S4). Since significant differences in *Symbiodinium* communities occurred between thermal regimes in *S. siderea* only, it is possible that nutrient loading or turbidity played a role in *Symbiodinium* variation within *S. siderea*, but may not have significantly influenced *Symbiodinium* communities in *S. radians* or *P. strigosa*. However, the magnitude of this influence cannot be teased apart from the effect of thermal regime without extensive quantification of nutrient concentrations across the Belize MBRS.

### Coral host may play a role in thermal tolerance

In this study, the relative abundance of thermally tolerant *Symbiodinium* D1a (*S. trenchii*) was not associated with inshore reefs as in Toller at al. [119], marginal reefs as in Hennige et al. [120] and LaJeunesse et al. [104], sites exposed to the highest temperatures as in Baker et al. [48], or sites exposed to the widest range of thermal fluctuations as in Abrego et al. [121], Fabricius et al. [122], and LaJeunesse et al. [40, 123]. Instead, *S. trenchii* was most prevalent at the southern Punta Gorda low_TP_ and mod_TP_ sites (Table S1, S5, Fig 3). Since *S. trenchii* is often associated with recently bleached and/or recovering corals [48, 124], but can be replaced or outcompeted following recovery [105], it is possible that a recent bleaching event may have occurred at these sites, however these data are not available. In summer 2014, temperatures at all sites in this study exceeded the published local bleaching threshold of 29.7°C [86] (Fig S1), yet *S. trenchii* was only the dominant symbiotic partner in eight *S. siderea* samples, all of which were from the same two sites (Punta Gorda low_TP_ and mod_TP_; Fig 3). The presence of *S. trenchii* in several *P. strigosa* corals taken from the Punta Gorda modTP site provides additional evidence of temperature stress at these sites (Punta Gorda low_TP_ and mod_TP_). This result suggests that corals at these sites had either bleached recently or retained *S. trenchii* as a dominant symbiont following past bleaching, possibly as a way to increase thermal tolerance [125]. Lower thermal tolerance has been proposed previously for *S. siderea [80]* and *Orbicella faveolata [126]* at these sites (Punta Gorda low_TP_ and mod_TP_) and may be due to nutrients, sediments, and low salinity terrestrial runoff exported from Guatemala and Honduras by currents that wash over this area of the Belize MBRS [126–128]. Low abundances of *S. trenchii* at other low_TP_ and mod_TP_ sites corroborates this hypothesis, as estimated thermal stress occurred at all latitudes at roughly the same magnitude (Fig S1). Overall, lack of *S. trenchii* in high_TP_ sites indicates that regardless of warmer and more variable conditions, these three coral species do not associate with this thermally tolerant symbiont. Therefore, presumed increased thermal tolerance at high_TP_ sites may be due to local adaptation of the coral host [37, 129] or strains of *Symbiodinium* [130, 131]. Further research into coral host and symbiont local adaptation would be needed to confirm this hypothesis.

## Conclusion

This study demonstrates that *Symbiodinium* communities associated with corals in Belize are dependent on both host species as well as environmental variables. *S. siderea Symbiodinium* communities were divergent from *S. radians* and *P. strigosa* (Fig 3; Fig 4A). Thermal regime played a role in driving *Symbiodinium* community composition in *S. siderea* but not *S. radians* or *P. strigosa*, suggesting that local impacts such as nutrients, sediment, or light availability may also influence *Symbiodinium* communities on the Belize MBRS. Additionally, low abundance of *S. trenchii* in inshore high_TP_ sites indicates thermal tolerance at these sites must be conferred through alternative mechanisms, such as local adaptation.

## Acknowledgements

We thank J. Watkins, L. Speare, and A. Knowlton for laboratory assistance and C. Berger for assistance with coding. We also thank NASA JPL and NOAA ERDAAP for access to MUR SST data used in this paper, Belize Fisheries Department for issuing research and collection permits, and Garbutt’s Marine for providing local expert guides and boats for field research. This work was supported by the Rufford Foundation (http://www.rufford.org) Small Grant to JHB (15802-1); National Science Foundation (Oceanography) (nsf.gov) to KDC (OCE 1459522); Department of Defense NDSEG fellowship to JHB. The authors declare that no conflict of interests exists.

